# Acetylcholine regulates pulmonary inflammation and facilitates the transition from active immunity to tissue repair during respiratory viral infection

**DOI:** 10.1101/2020.07.02.184226

**Authors:** Alexander P. Horkowitz, Ashley V. Schwartz, Carlos A. Alvarez, Edgar B. Herrera, Marilyn L. Thoman, Dale A. Chatfield, Kent G. Osborn, Ralph Feuer, Uduak Z. George, Joy A. Phillips

## Abstract

Inflammatory control is critical to recovery from respiratory viral infection. Acetylcholine (ACh) secreted from non-neuronal sources, including lymphocytes, plays an important, albeit underappreciated, role in regulating immune-mediated inflammation. This study was designed to explore the role of ACh in acute viral infection and recovery. Using the murine model of influenza A, cholinergic status in the lungs and airway was examined over the course of infection and recovery. The results showed that airway ACh remained constant through the early stage of infection and increased during the peak of the acquired immune response. As the concentration of ACh increased, cholinergic lymphocytes appeared in the airway and lungs. Cholinergic capacity was found primarily in CD4 T cells, but also in B cells and CD8 T cells. The cholinergic CD4+ T cells bound to influenza-specific tetramers at the same frequency as their conventional (i.e., non-cholinergic) counterparts. In addition, they were retained in the lungs throughout the recovery phase and could still be detected in the resident memory regions of the lung up to two months after infection. Histologically, cholinergic lymphocytes were found in direct physical contact with activated macrophages throughout the lung. When ACh production was inhibited, mice exhibited increased tissue inflammation, altered lung architecture, and delayed recovery. Together, these findings point to a previously unrecognized role for ACh in the transition from active immunity to recovery and pulmonary repair following respiratory viral infection.

## Introduction

A growing body of research indicates that acetylcholine (ACh) produced by specialized lymphocytes plays a critical role in regulating inflammation and immunity(1-6). It is now understood that roughly 60% of the ACh content of whole blood is sequestered within mononuclear leukocytes (MNLs), a group comprised of mostly lymphocytes and a small number of monocytes(5-8). Immune cells preform and store ACh in a similar fashion to neurons(9), allowing for quick and efficient release following appropriate stimulation. Modulation of immune inflammatory responses by ACh occurs in a site and target specific manner(10). Vagal nerve derived norepinephrine (NE) induces ACh release from cholinergic CD4^+^ T cells in the spleen, whereas cholinergic B-1 cells in the peritoneal cavity release ACh in response to TLR agonists, surface Ig ligation, and cholecystokinin (CCK) (11, 12). During chronic hepatitis, both CD4 and CD8 cells secrete ACh in response to IL-21 (13).

ACh decreases inflammation in large part by action on local macrophages. Signal transduction via the alpha-7 nicotinic ACh receptor (α7-nAChR) decreases nuclear translocation of the Nf-κB transcription factor, ultimately reducing macrophage production of the pro-inflammatory cytokine TNF (10, 14). Ligation of α7-nAChR significantly reduces lung injury and overall mortality in septic shock models (14, 15). In addition, pro-inflammatory cytokines TNF, IFN-γ, and IL-6 are increased in mice lacking the α7-nAChR, further indicating that α7-nAChRs play a critical role in regulating macrophage-associated inflammation (16).

Efficient inflammatory regulation is a crucial feature of recovery from respiratory viral infection (17). Overwhelming inflammation during viral illness, commonly called a “cytokine storm” results in significant lung damage and greatly increased risk of death. This is well established for influenza as well as the novel coronavirus COVID-19 (18, 19). Conversely, the decreased ability to properly induce and regulate inflammation displayed by the elderly is also associated with increased overall morbidity and mortality from respiratory infection (19-21).

This study was designed to explore the role of ACh during respiratory infection. Our results show that the airway ACh concentration changes over the course of infection, and that the changes are mirrored by an influx of cholinergic lymphocytes. Inhibiting ACh synthesis resulted in extended pulmonary inflammation, increased macrophage activation, and delayed tissue repair. These findings illuminate a previously unknown role of ACh in recovery from acute viral infection, and further illuminate the non-neuronal cholinergic system as an underappreciated therapeutic target for inflammatory regulation, particularly during viral infection (22-24).

## Methods

### Ethics Statement

All animal experimental protocols were approved by the Institutional Animal Care and Use Committee at San Diego State University (protocol numbers 15-06-006) and 18-06-008P) prior to initiation of experiments. Animals were given free access to food and water at all times and were cared for according to guidelines set by the American Association for Laboratory Animal Care.

### Animal Experimentation

Conventional C57BL/6 and ChAT (BAC) – eGFP transgenic mice [B6.Cg-Tg(RP23-268L19-EGFP)2Mik/J] with endogenous choline acetyltransferase (ChAT) transcriptional regulatory elements directing eGFP expression were originally obtained from The Jackson Laboratory (JAX, Bar Harbor, ME) and were bred in house. Animals were used between the ages of aged 12-20 weeks. Mice were infected with the mouse adapted influenza virus A/Puerto Rico/8/34 (H1N1) (PR8) (Charles River Laboratories Avian Vaccine Services, North Franklin, CT) delivered in a volume of 30 □l exactly as in(25). All animals were examined for overt indications of morbidity and weighed daily starting the day of infection. For euthanasia, animals were exposed to isofluorane until respiration ceased. Death was confirmed by severing the abdominal aorta. For inhibition of ACh synthesis, Hemicholinium-3 (HC3) (Millipore Sigma, Burlington, MA) was administered via intraperitoneal (IP) injection. Animals were treated daily for six days, beginning seven days post infection (dpi). Each animal was administered 100 μL of a 10 μg/mL HC3 solution dissolved in PBS for a final dose of 1 μg of HC3 daily. Control animals were administered 100 μL of sterile PBS by IP injection on the same days.

### Flow Cytometry

Following euthanasia, lungs were lavaged with 1 mL of sterile PBS as in in [Sanderson, 2012 #696]. Airway cell counts were immediately collected using an Accuri C6 flow cytometer (Accuri Cytometers, Ann Arbor, MI) prior to centrifugation and separation of the airway cell pellet from the cell free bronchial alveolar lavage (BAL) fluid. Lungs were removed, processed, and stained for flow cytometry as in [Sanderson, 2012 #696]. Flow cytometry antibodies used in this study were: 1A8-FITC (BD PharMingen, San Jose, CA), CD4-APC,, CD8-PE/Cy5, B220-PE (eBioScience, San Diego, CA), CD11b-APC (BioLegend, San Diego, CA), CD11c-PE/Cy5 (Tonbo Bioscience, San Diego, CA). To examine influenza specificity, lung and airway lymphocytes were stained using the influenza-specific CD4 tetramer: I-A(b) Influenza A NP 311-325 QVYSLIRPNENPAHK as described(26). Negative control CD4 tetramer was I-A(b) human CLIP 87-101 PVSKMRMATPLLMQA. Tetramers were obtained from the NIH Tetramer Core Facility. Samples were incubated with tetramer for 60 minutes at room temperature. For all flow cytometric studies, data acquisition and analysis were performed on an Accuri C6 (BD Biosciences, San Jose, CA) flow cytometer using CFlow Plus Analysis software.

### Mass Spectrometry

Choline and ACh were measured in cell-free BAL fluid by hydrophilic interaction liquid chromatography coupled to tandem mass spectrometry (HILIC LC–MS/MS), using stable isotope-labeled internal standards (choline-d4 and acetylcholine-d4) as described (27, 28). Briefly, an aliquot of the cell-free BAL fluid was spiked with a mixture of deuterated choline/deuterated ACh. The sample was adjusted to 50% methanol and ice partitioned to remove proteins. The remaining sample was lyophilized, dissolved in acetyonitrile:H20 and analyzed using a Thermo-Finnigan TSQ Quantum LC-MS/MS in positive ion ESI mode. For quantification, MS/MS ion transitions m/z 104 to 60 (choline), 108 to 60 (choline -d4), m/z 146 to 87 (ACh), and 150 to 91(ACh-d4) were used.

### Histology

Following euthanasia, exsanguination, and cannulation of the trachea, lungs were inflated with 1mL of a 4% paraformaldehyde solution. The trachea was clamped and all lungs were removed and fully submerged in the 4% paraformaldehyde solution overnight fixation (~18 hour), with hemostats still attached and fully restricting flow through the trachea. The following day, hemostats were removed, lung lobes were dissected from one another and fully submerged in 70% Ethanol (EtOH), which was replaced 6-8 hours. One day prior to processing, EtOH was discarded and replaced for a further 24 hours before being sent for processing and paraffin embedding. Formalin fixed paraffin embedded (FFPE) lung lobes were serially sectioned at 5um on a Leica RM2125 RTS Rotary Microtome, and mounted on Prism (Prism Research Glass, Raleigh, NC) positively charged microscope slides. Formalin fixed paraffin-embedded (FFPE) sections were stained with Hematoxylin and Eosin (H&E) for histological analysis using standard methods. Tissue sections of uninfected, infected vehicle control, and infected HC3 treated lung tissue were analyzed by a veterinary pathologist who was blinded to treatments and groups.

For immunofluorescence studies, FFPE sections were deparaffinized and treated for immunofluorescent imaging by standardized methods(29). Sections were blocked with a 10% normal goat serum (NGS) solution for 30 minutes and incubated with primary antibody overnight at 4°C. All antibody dilutions were made in a 2% normal goat serum solution. Detection of GFP and Iba1 required high-temperature antigen unmasking in 0.01 M citrate buffer (pH 6.0) (Sigma-Aldrich, San Diego, CA). Sections were treated with biotin and streptavidin blocking solutions (Vector Laboratories, Burlingame, CA) before incubation with primary antibodies. Sections were incubated with a rabbit primary monoclonal antibody against Iba1 [ionized calcium-binding adapter molecule 1 (Iba1): polyclonal rabbit anti-Iba1 antibody; Wako Pure Chemicals Industries, Ltd, Osaka Japan] at 1:500 at 4C overnight. A goat secondary antibody [goat biotinylated anti-rabbit IgG (H+L); Vector Laboratories, Burlingame, CA] at 1:500 was diluted in 2% normal goat serum and incubated on sections for 30 minutes. After staining with secondary antibodies, all sections were washed three times with phosphate buffered saline and incubated with a streptavidin-AlexaFluor 594 complex [DyLight 594 streptavidin conjugate; Vector Laboratories, Burlingame, CA] 1:500 diluted in 2% normal goat serum. For tri-color stained sections, sections were incubated for 30 minutes in a light free environment with an anti-GFP AlexaFluor-488 conjugate [anti-GFP, rabbit polyclonal antibody, Alexa Fluor 488 conjugate; Invitrogen, Eugene, OR] diluted 1:5 in 2% normal goat serum. Specificity controls for immunostaining included sections stained in the absence of primary antibody or in the presence of rabbit immunoglobulin G control antibody at 0.1 ug/mL (Vector Laboratories, Inc.). Sections were overlaid with Vectashield anti-fade mounting medium (Vector Laboratories, Burlingame, CA) containing DAPI (4’, 6-diamidino-2phenylindole) to detect DNA/nuclei (blue) and covered with glass coverslips. Sections were observed by fluorescence microscopy (Ziess Axio Observer D1 Inverted Phase Contrast Fluorescent Microscope). Green, red, and blue channel images were merged using AxioVision software. Broad field images were taken on a Zeiss Axio Zoom.V16 Stereo Zoom Microscope (Zeiss, Germany).

### Automated Image Segmentation and Quantification of Immunofluorescence

For regional comparison of the lungs between treatment groups, images were categorized into three categories based on anatomical region: open alveolar space (open), bronchus associated lymphoid tissue (BALT), and area peripheral to large airways (peri-bronchial). All images to be quantified were captured on a Ziess Axio Observer D1 Inverted Phase Contrast Fluorescent Microscope at the same magnification, utilizing the 20x objective with a digital zoom of 0.97. Exposure times were held constant in red, green, and blue channels for each image captured, although only the signal in the red channel was quantified, as Iba1 was marked with the AlexaFluor 594 fluorochrome. Exposure times were: red (2.1s), green (2.1s), blue (200ms). The same exposure time was utilized for both the red and green channels for comparison of auto fluorescent tissues within each section, such as red blood cells and fibrin deposits, commonly seen as a result of vascular leakage in inflamed tissues. For Iba1 immunofluorescence quantification, only images in the red channel were analyzed.

### Automated Image Segmentation

A novel automated algorithm was designed and implemented in MATLAB 2019b Image Processing Toolbox to accurately quantifying the red stains present in the lungs. The program locates and segments regions of interest while simultaneously calculating the size and intensity of these regions. With a goal of capturing the red stain present in the images, the gray-scale red channel portion of the original microscopy images were used for the automated image analysis process. The program follows three main steps: image input, image segmentation, and data extraction. The step for the image input was automated for efficiency by directing the program to read multiple images in a folder sequentially. After an image is read, the program segments the red dots using Otsu’s thresholding method in which background noise in the image is eliminated by selecting a threshold automatically from a gray level histogram using discriminant analysis(30). While there are a variety of thresholding methods present in the literature, Otsu’s method is the most accurate and most widely used(31-34). In the program, Otsu’s method determines a threshold that distinguishes the background from the region of interest. The determined threshold is then used to segment each image, removing background noise and displaying the red stain. The final step is the calculation of the total area of red stain present in the image as well as the intensity of the stains. Total pixel area of red stain coverage is calculated by determining the number of nonzero elements in the gray scale image while total image intensity is calculated by summing the intensity levels of all elements remaining in the segmented image. The program analyzed all the microscopy images and generated graphs for total area of red dots and intensity in approximately 29.6 seconds.

### Statistical Analysis

Statistical analysis was computed using R and R Studio. One-way ANOVA and two-tailed paired t-tests were performed *, p<0.05 was considered statistically significant.

## Results

### Airway choline changes during infection and recovery

Animals were infected with a sublethal dose of influenza A/PR8 (H1N1) and weighed daily to monitor morbidity (Fig 1A). Separate cohorts were euthanized at specified time points and BAL fluid was isolated in order to measure the airway ACh concentration. Airway choline concentration was also measured as a biomarker for local ACh hydrolysis(23). There was no change in the airway ACh concentration over time; however, the airway choline concentration changed over the course of infection. From a background concentration of 1200ng/ml prior to infection, the airway choline concentration increased to 4800 ng/mL 8 dpi and peaked at 6500 ng/mL 10 dpi. By 15 dpi, BAL choline concentration diminished to 1800 ng/mL, similar to the starting concentration measured 0 dpi (Fig 1B). Comparing the weight change curve to the choline concentration changes shows that ACh hydrolysis reaches a peak in the influenza-infected lungs shortly after the point of peak weight loss.

To determine the source of airway ACh and examine a possible role for non-neuronal ACh production during influenza infection, we examined the kinetics of lymphocyte populations infiltrating airways and lungs during infection using ChAT (BAC) – eGFP transgenic mice (ChAT mice) with endogenous choline acetyltransferase transcriptional regulatory elements directing eGFP expression, alongside C57BL/6 mice as non-reporter controls. Wild-type, influenza-infected C57BL/6 mice were used as controls for green fluorescent protein (GFP) fluorescence. Starting 8 dpi, GFP^+^ cells were detected in the airway of the ChAT mice (Fig 1C). The number of ChAT-GFP^+^ lymphocytes present in BAL samples followed the same kinetic pattern as the total lymphocyte population (Fig 1D), increasing rapidly starting 7 dpi and reaching peak numbers between 8-10 dpi (Fig 1E). However, the percentage of ChAT-GFP^+^ lymphocytes remained above 7% of the total lymphocyte population in the airways through 28 dpi (Fig 1F).

**Figure 1:**
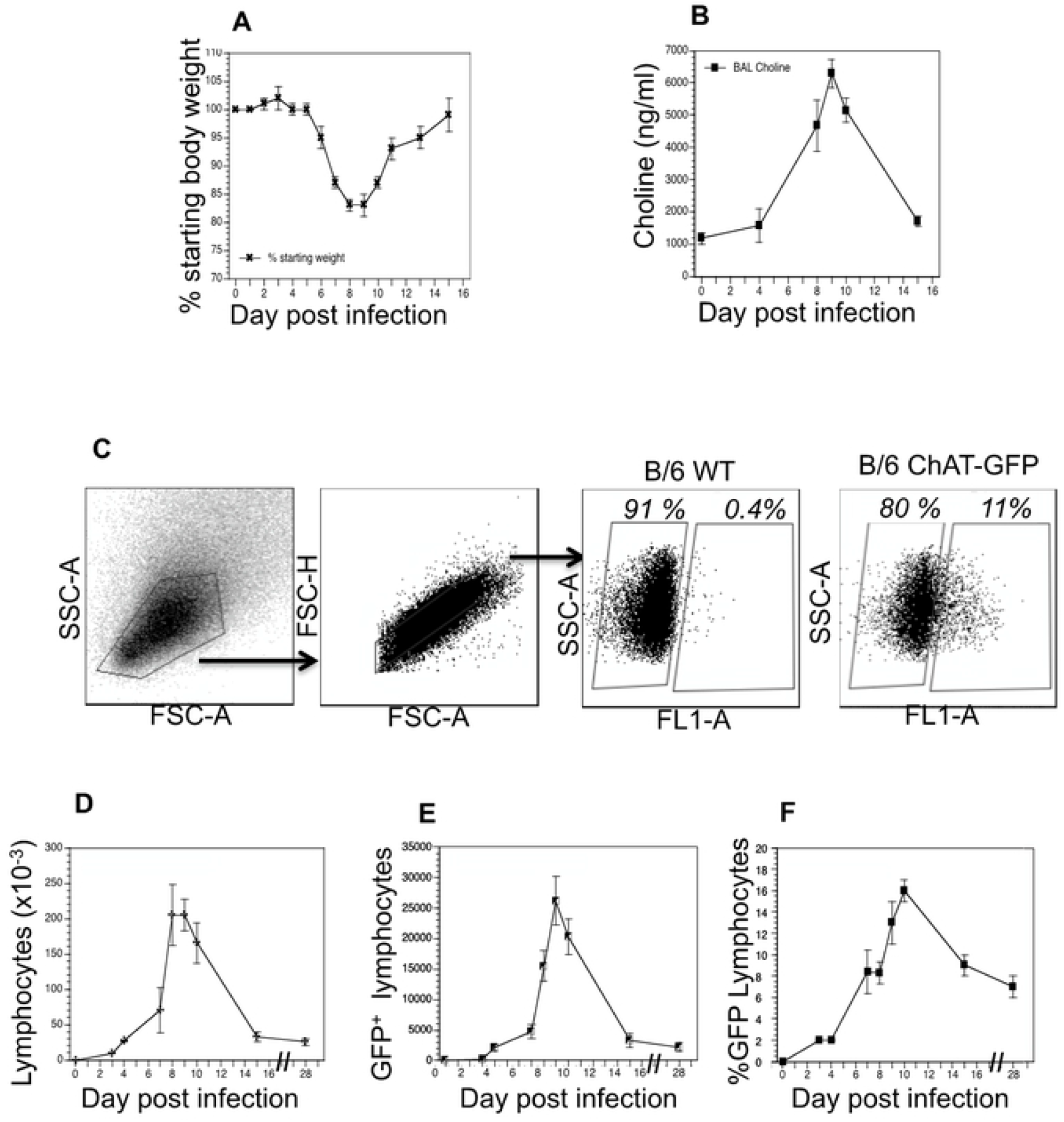
Cholinergic status kinetics in the influenza-infected Lung. Mice were infected with a non-lethal dose of influenza (0.336MLD50) as described in Materials and Methods. Mice were weighed daily starting the day of infection. **A.** Weight change following influenza infection. Group average weight change is shown as mean ± SEM. **B.** Airway choline concentration was measured using HILIC LC–MS/MS with a stable choline-d4 isotope-labeled internal standard as described in Materials and Methods. **C**. Ten dpi, airway cells were isolated by lung lavage, stained with fluorescent antibodies and analyzed by FACS as described in Materials and Methods. FACS plots shown are from individual mice representative of at least 10 mice per time point. Gating strategy for BAL cell analysis is shown using wild-type C57Bl/6 mouse as a negative control for GFP fluorescence. Representative staining data is shown (10–30 mice per time point). FSC: forward scatter (size); SSC: side scatter. Cholinergic capacity was defined as increased FL1 fluorescence compared to wild-type C57Bl/6. **D**. Kinetics of total airway lymphocytes; **E**. Kinetics of airway cholinergic (GFP^+^) cells; **F**. Percentage of all lymphocytes expressing GFP over the course of influenza infection.

Flow cytometry identified ChAT-GFP^+^ subsets of both CD4^+^ and CD8^+^ T cell populations in the airways during peak days of infection (8-10 dpi) (Fig 2). The majority of these ChAT-GFP^+^ lymphocytes were CD4^+^ (69.7%) while 23.3% were CD8^+^, indicating more cholinergic helper T cells present in BALF samples than cholinergic cytotoxic T cells, respectively. When the analysis was extended to lung tissue, ChAT-GFP was detected in B220^+^ B-1 lymphocytes as well as CD4 and CD8 T cells. In airway and lung tissue, the highest percentage of GFP^+^ cells were CD4^+^ T cells. The CD4 population also expressed the most GFP fluorescence on a per-cell basis (Table 1).

**Figure 2.**
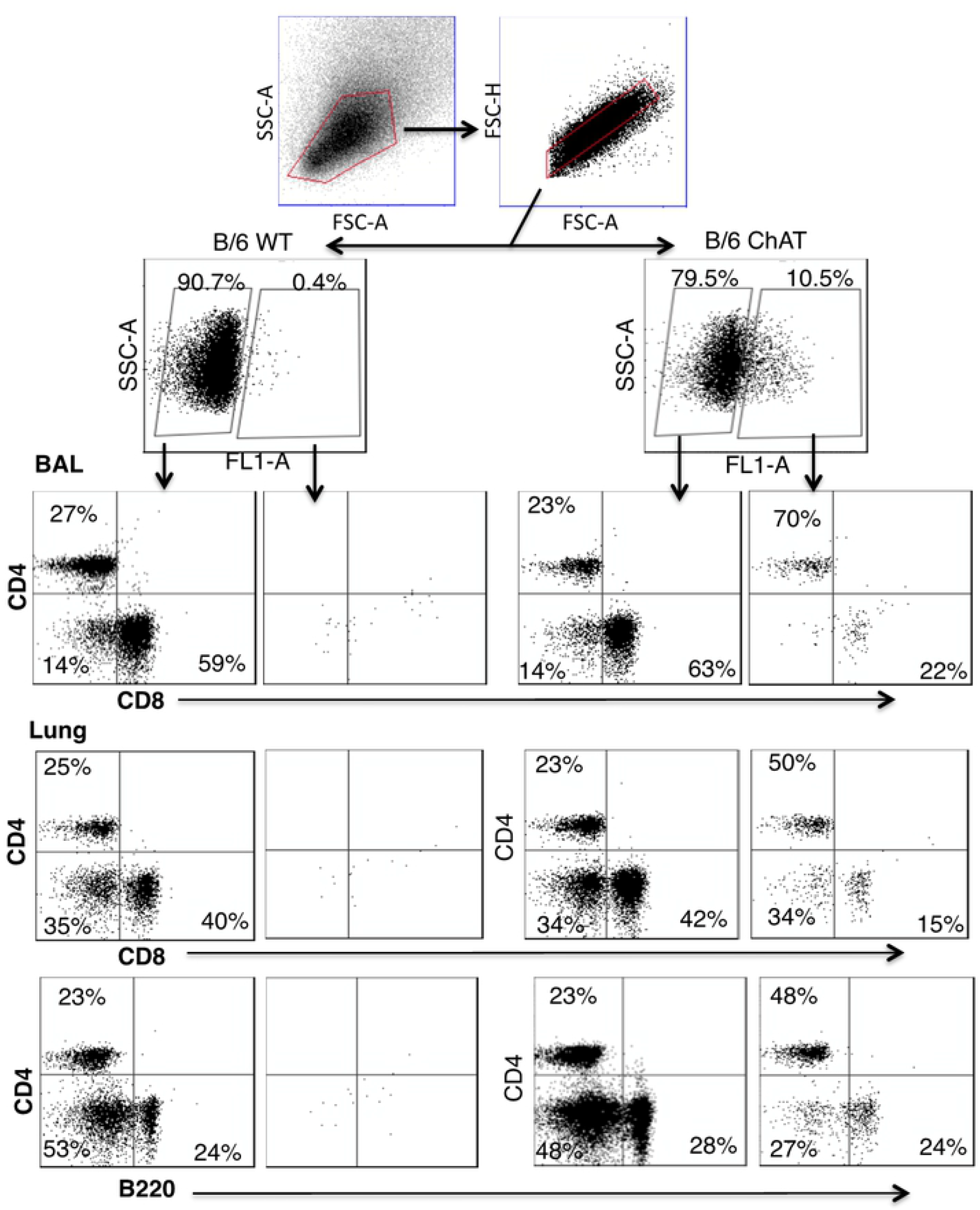
Airway and Lung Cholinergic Cell Phenotyping. Animals were infected with influenza and sacrificed ten days later. Cells were isolated and stained for surface marker analysis as described in Materials and Methods. Gating strategy to examine surface phenotypes of FL-1^+^ and FL-1^-^ cells is shown.

**Table 1.** Pulmonary Cholinergic Lymphocyte Phenotype. Day 10 post infection. Data compiled from five experiments, mean ± SE

|  | % ChAT-GFP | ChAT-GFP MFI |
| --- | --- | --- |
| BAL CD4 | 26 + 1 | 21423 + 2127 |
| Lung CD4 | 21 + 2 | 21278 + 2110 |
| BAL CD8 | 6 + 2 | 7893 + 3222 |
| Lung CD8 | 4 + 1 | 9972 + 3324 |
| Lung B220 | 11 + 2 | 13749 + 1393 |

Influenza antigen specificity of the TCR in cholinergic and conventional T cell populations was examined by tetramer staining (Fig 3). Neither conventional nor cholinergic CD4 T cells bound to fluorescent tetramers loaded with negative control peptide (human CLIP 87-101). Similar percentages of cholinergic CD4 T cells and conventional T cells bound the dominant influenza A epitope NP311-325 (Fig 3).

**Figure 3.**
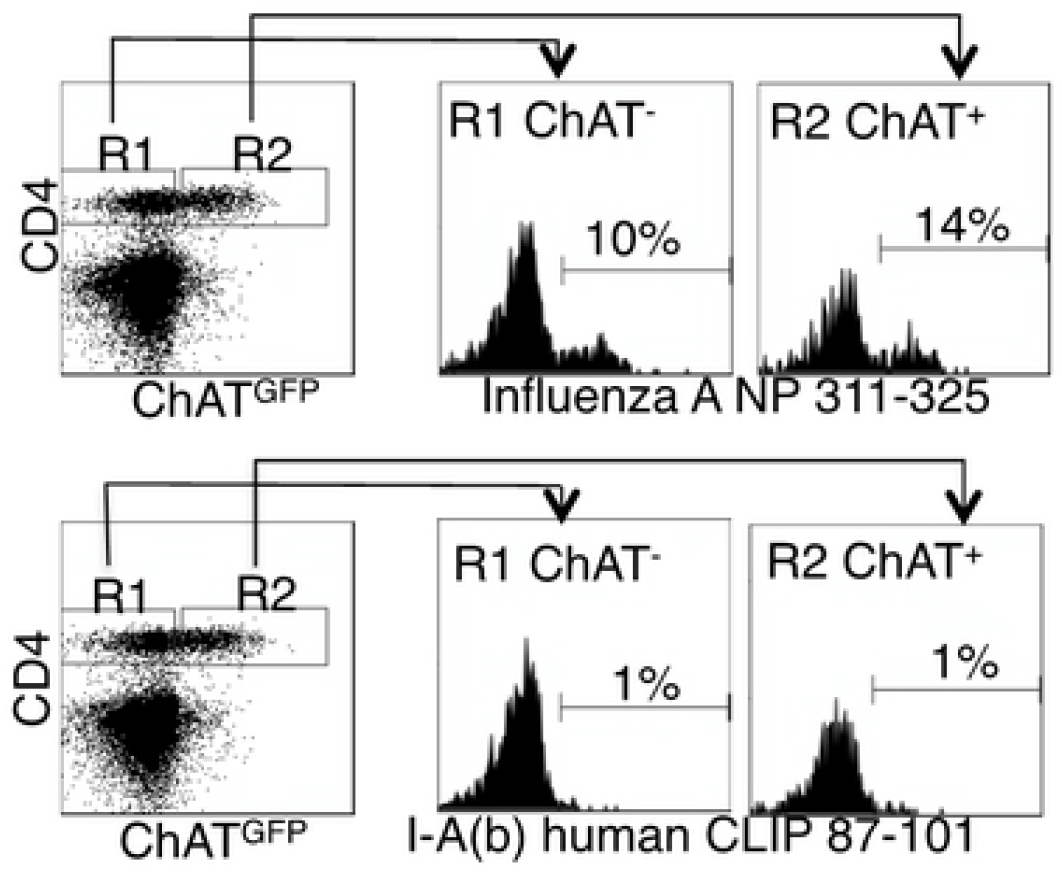
Cholinergic CD4 T cells bind to influenza-specific tetramers. Mice were infected with influenza A/PR8 and sacrificed for analysis 8 days later. BAL cells were isolated and stained the anti-CD4 as in figure 2. Cells were then stained with either the class II tetramer I-A(b) Influenza A NP 311-325 (top) or the negative control tetramer I-A(b) human CLIP 87-101 (bottom). Histograms show staining of either R1 = conventional CD4 T cells; R2 = cholinergic CD4 T cells vs. ChAT-GFP fluorescence.

Surface staining indicated that the cholinergic CD4 T cells were uniformly CD44^hi^CD62U^lo^ (Fig 4). This matches the overall surface phenotype of cholinergic CD4 T cells from multiple reports, but it also corresponds with a specific subset associated with long term memory known as the T resident memory population. To explore the possibility that cholinergic CD4 T cells made up part of the TRM population, mice were infected with influenza A and allowed to recover for two months with no manipulation, then they were given an intravenous injection of fluorescent anti-CD45 antibody and sacrificed ten minutes later. As shown in Fig 4B, CD4 positive cells in the lungs can be divided into two subsets, based on staining by the injected CD45. Those cells not exposed to the circulation were left unstained following iv injection. These represent the T resident memory (TRM) population. The T effector memory (TEM) reside in areas of the lung accessible to the circulation and were stained following iv injection of fluorescent antibody(35). Cholinergic CD4 T cells were primarily found in the regions of the lung sequestered from circulation, associated with the TRM population (Fig 4B).

**Figure 4.**
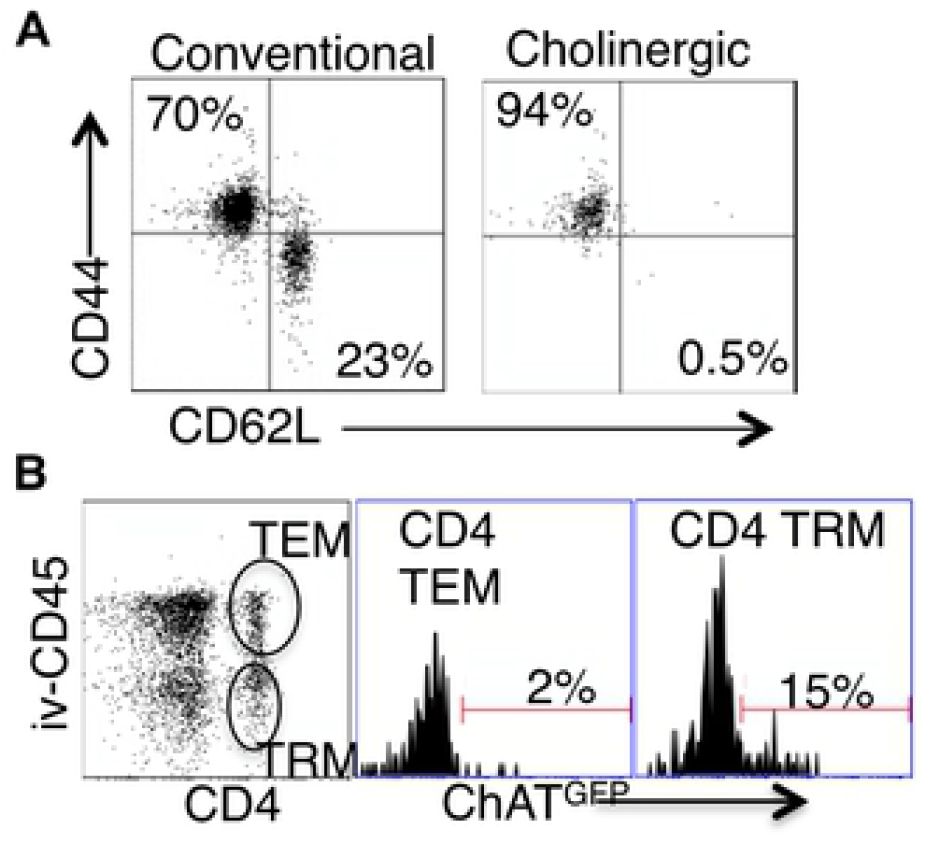
Cholinergic CD4 T cells reside in the resident memory niche of the lung. Two months after influenza infection, mice were injected intravenously with fluorescent anti-CD45 ten minutes before sacrifice. Lung lymphocytes were stained for surface markers and analyzed based on GFP expression as in Figures 1 and 2. **A.** Gated CD4 cells were stained for memory markers CD44 and CD62L. **B.** Total lymphocytes from ChAT-GFP mice were analyzed based on fluorescence of the infected CD45 antibody vs CD4 expression. CD4 positive populations were then analyzed for GFP expression.

### Cholinergic Lymphocytes associate with activated macrophages

To explore the localization of cholinergic cells within the context of the overall lung architecture, lungs from animals inflected with influenza A were fixed and processed for immunofluorescent (IF) staining and analysis. Cholinergic cells were identified by staining with an anti-GFP fluorescent antibody. The cholinergic GFP^+^ cells were predominantly localized to bronchus associated lymphoid tissues (BALT) but they were also observed in open spaces, specifically the alveolar space and peri-bronchial regions proximal to the major sites of active inflammation or infection, identified by Iba1 staining. Iba1, also known as allograft inflammatory factor 1 or AIF-1, is a marker of activated macrophages and ongoing inflammation(8, 29, 36-38). As shown in Fig 5, cholinergic lymphocytes were regularly observed in direct physical contact or close spatial proximity with Iba1^+^ activated macrophages. Co-expression of ChAT-GFP and Iba1 was never observed within the same cell in vehicle control or HC3 treated lungs.

**Figure 5.**
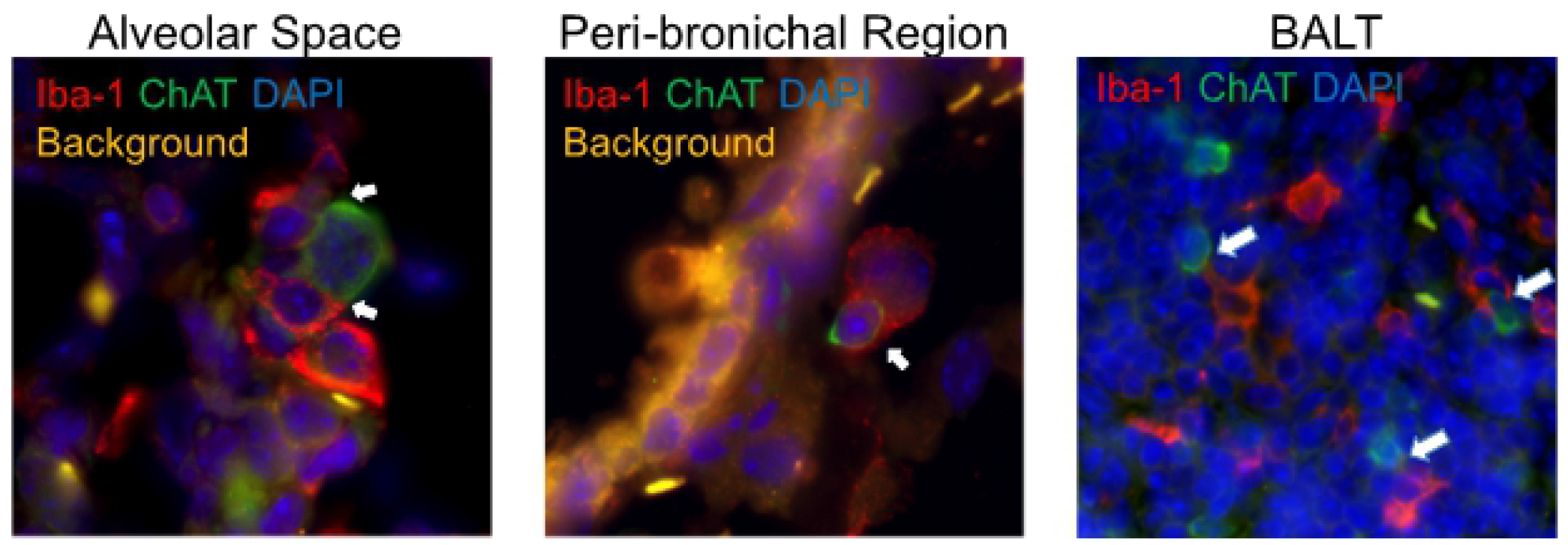
Cholinergic lymphocytes are found in direct contact with activated macrophages throughout the lung. Dual labeled sections (Green: ChAT-GFP, Red: Iba1) of infected vehicle control and HC3 treated lungs from 12 dpi show close contact of cholinergic lymphocytes (ChAT-GFP^+^) and activated macrophages (Iba1^+^). Alveolar space and Peri-bronchial region images captured at 63x oil immersion with 1.4x digital zoom. BALT image captured at 40x oil immersion with 1.3x digital zoom. Arrows point out areas of lymphocyte-macrophage contact.

### Blocking Ach synthesis increases viral-associated morbidity

The co-localization of cholinergic lymphocytes and activated macrophages indicated a potential role for targeted ACh delivery to activated macrophages during the later stages of influenza infection. To examine the requirement for ACh during recovery from influenza infection, we used the choline reuptake inhibitor Hemicholinium-3 (HC3) to disrupt ACh synthesis during the time period associated with the increased airway ACh concentration (Fig 1). Mice were infected with influenza A/PR8 (H1N1) as above. After seven days, mice were stratified into treatment cohorts according to the amount of weight loss to ensure equivalent pre-established morbidity in each cohort. One cohort was treated with HC3 from days 7 through 12 (infected HC3 treated), while the infected vehicle control cohort was injected with saline to control for handling/injection stress. Additional control cohorts were injected with HC3 or PBS without having been infected. One infection cohort was sacrificed 10 dpi to measure the airway choline concentration. BAL samples from the HC3 treated cohort exhibited increased airway choline concentration compared to infected vehicle control groups, indicating that HC3 was inhibiting choline uptake in the pulmonary airways (Fig 6A). All influenza infected cohorts lost weight as expected. The infected control cohort began to regain weight 9 dpi and had returned to 96% of their starting weight by 15 dpi. In contrast, the infected HC3 treated cohort did not begin to regain weight until 11 dpi and only returned to 91% of their starting weight by 15 dpi. The uninfected cohort treated with HC3 did not display any treatment-associated weight change (Fig 6B). Flow cytometric analysis showed an increase in the number of neutrophils in HC3 treated animals compared to vehicle control animals (Fig 6C).

**Figure 6.**
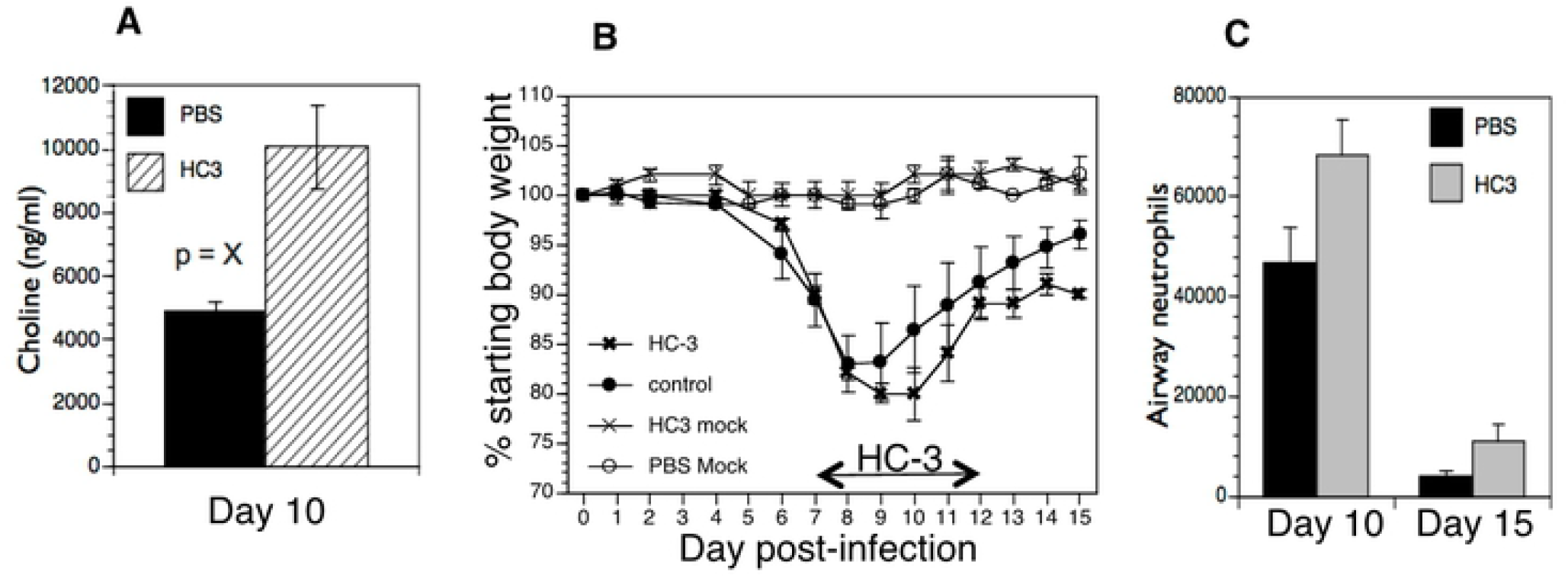
Decreasing ACh synthesis delays recovery and increases inflammation following influenza infection. Mice were infected with influenza and injected with HC3 or PBS 7-12 dpi as described in Materials and Methods. **A**. Airway choline was measured ten days after infection as in Figure 1. **B**. Weight was measured daily throughout infection and recovery. Control cohorts were injected with either HC3 or PBS but were not infected with influenza. Group average weight changes are shown as mean ± SEM. **C**. On days 10 and 15 after infection, pulmonary neutrophils (SSC^hi^CD11b^+^Ly6G/1A8^+^) in the influenza-infected cohorts were identified by FACS analysis.

### Blocking Ach synthesis increases pulmonary inflammation

We used Iba1 as a marker of overall inflammation(8, 29, 36-38) to determine the effect of HC3 treatment during the later stage of infection (Fig 7). Fluorescent signal from immunofluorescence stained slides was quantified spatially to determine the degree of inflammation at set time points. A novel automated algorithm was designed and implemented in MATLAB 2019b Image Processing Toolbox to accurately quantifying the red stains present in the lungs. The program locates and segments regions of interest while simultaneously calculating the size and intensity of these regions, using Otsu’s method of thresholding in order to remove background noise and isolate the red stain with minimal human bias(30, 32-34). Area of stain, which corresponds to the total number of pixels occupied by fluorescent signal from any one image, was evaluated by region and compared between infected HC3 treated groups and infected vehicle control groups. Area of stain from Iba1 fluorescence differed between regions of the lung in both HC3 treated animals and vehicle control animals (Fig 7B). Area of stain measurements from infected HC3 treated groups were greater than those observed in infected vehicle control groups in all three lung regions, as well as tissue wide. Total intensity, corresponding to the sum of fluorescent stain intensity per pixel in one image, was measured by an novel automated image segmentation algorithm and evaluated by region and compared between influenza-infected HC3 treated vs. vehicle control groups (Fig 7C). Total intensity was determined to be greater in infected HC3 treated groups compared to infected vehicle control groups in all three lung regions, as well as tissue wide.

**Figure 7:**
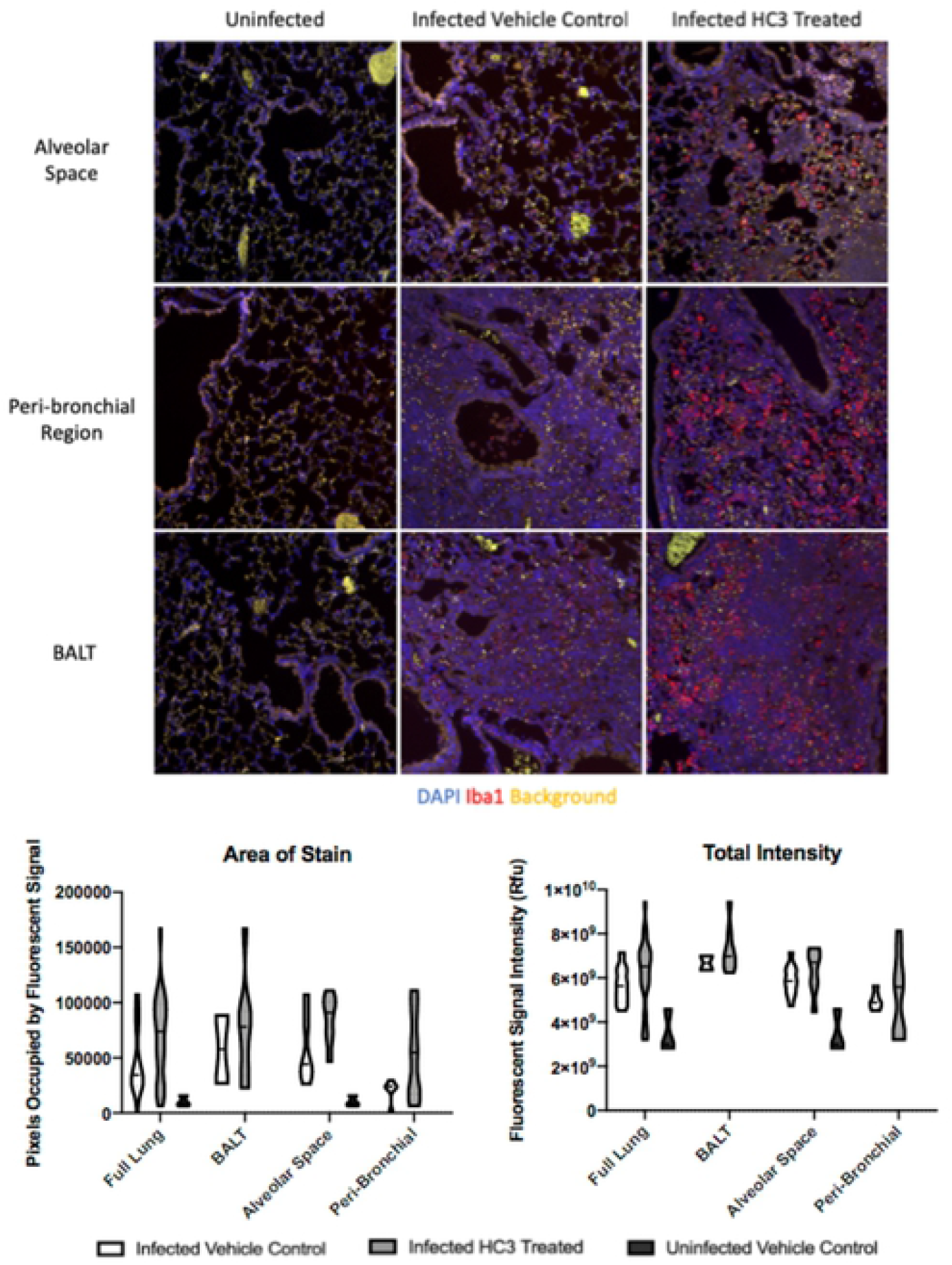
IHC Analysis of Iba1 During Recovery. A. Immunofluorescence of Iba1 staining in multiple lung regions 15dpi. Representative images taken at 20x showing Iba1 staining (red, AlexaFluor-594) and DAPI stained nuclei (blue) in the alveolar space, peri-bronchial region, and bronchus associated lymphoid tissues (BALT) in uninfected, infected vehicle control, and infected drug treated animals 15 dpi. Increased intensity and area of stain was observed in all lung regions of infected HC3 treated groups when compared to infected vehicle control groups. B. Quantification of Iba1 staining in HC3 and Control animal immunofluorescence sections. Area of stain analysis indicated more numerous pixels occupied by fluorescent signal in the lungs of infected HC3 treated groups compared to infected vehicle control groups in all three lung regions individually. Area of stain was also greater in infected HC3 treated groups compared to infected vehicle control groups when measurements of three lung regions were combined into a single data set. C. Total intensity analysis identified greater total intensity of fluorescent signal in the lungs of infected HC3 treated groups compared to infected vehicle control groups in the alveolar space and peri-bronchial regions, as well as the full lung when all measurements were compiled into a single data set. Greater total fluorescent intensity was also observed in the BALT region of infected HC3 treated groups compared to infected vehicle control groups, but was less significant than other regions or tissue wide.

### Decreasing Ach results in increased lung pathology

Lung samples from the HC3-treated animals and controls were collected 15 dpi, fixed and processed for staining examined to determine the effect of ACh disruption on tissue repair. As shown in Fig 8, infected vehicle control lung tissue was characterized by having mild/moderate multifocal perivascular mixed infiltrate and alveolar/interstitial mixed infiltrate of lymphocytes and neutrophils. However, infected HC3 treated lung tissue exhibited histological abnormalities not seen in the vehicle control lungs. The lungs from infected animals treated with HC3 were characterized as having moderate perivascular as well as alveolar/interstitial mixed infiltrate of lymphocytes and neutrophils (Fig 8A). In addition, pathological anomalies including as multifocal type II pneumocyte proliferation (Fig 8B), mild multifocal squamous cell metaplasia (Fig 8C), and mild fibroplasia (Fig 8D) were observed in infected HC3 treated lung tissue but were absent in infected vehicle control lung tissue.

**Figure 8:**
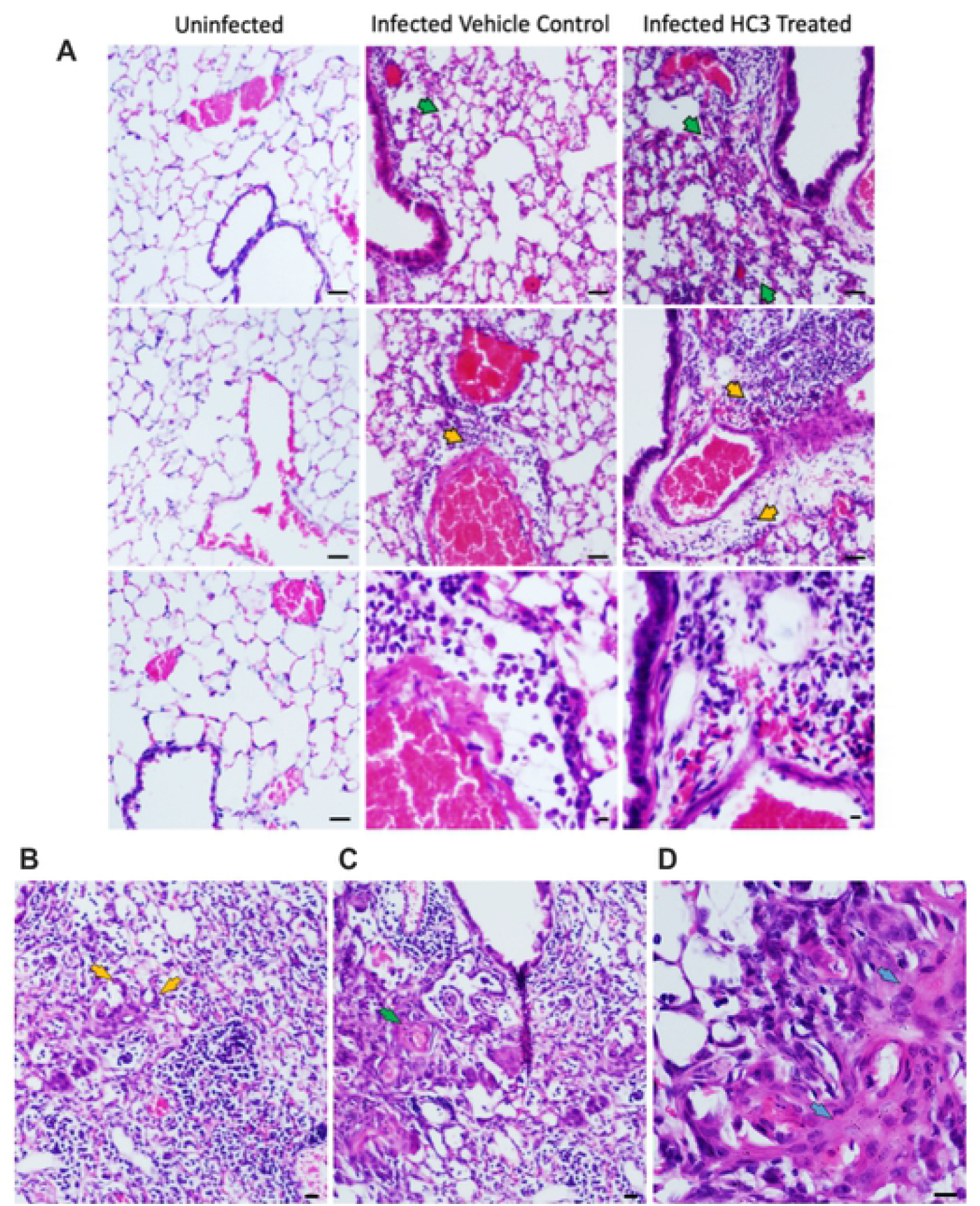
Histological Analysis of Recovery. A. Representative images of H&E stained sections of lung tissue at 10x (scale bar 100um) and 40x (scale bar 10um) magnification from healthy animals, infected vehicle control animals, and infected HC3 treated animals from left to right. Flu infected vehicle control lung tissue with mild/moderate multifocal perivascular mixed infiltrate (yellow arrow) and alveolar/interstitial mixed infiltrate of lymphocytes and neutrophils (green arrow). Flu infected HC3 treated lung tissue with moderate perivascular (yellow arrows) as well as alveolar/interstitial mixed infiltrate of lymphocytes and neutrophils (green arrows). B-D. Infected HC3 treated lung tissue shows multifocal type 2 pneumocyte proliferation (yellow arrows), mild multifocal squamous cell metaplasia (green arrows), and mild fibroplasia (blue arrows).

## Discussion

The present study adds to a growing body of research illuminating the crucial role of non-neuronal ACh in regulating inflammation and immunity. Our results demonstrate that ACh plays an important role in recovery from pulmonary viral infection. Inflammatory control and pulmonary tissue repair is critically important during respiratory infection, especially in the alveolar space where the majority of gas exchange occurs. Disruption of the delicate architecture of the alveolar space by direct viral damage and inflammatory cell influx during infection results in significant reduced tidal volume and a decreased capacity for gas exchange.

These studies used influenza A; however, the results are likely to be applicable to many acute respiratory infections including the pandemic SARS-CoV coronavirus COVID-19. As widely observed during the yearly influenza season as well as the current COVID-19 pandemic, many patients in critical condition exhibit significantly reduced tidal volume and capacity for gas exchange, resulting in the critical need for ventilators worldwide. Our findings that ACh from cholinergic lymphocytes regulates pulmonary inflammation and plays a role in tissue repair point to a previously unexploited therapeutic target for treating respiratory infection.

Based on previously published studies demonstrating ACh regulating inflammatory cytokine expression, macrophage activation, and neutrophil trafficking(11, 12, 14, 39-43), we initially hypothesized that ACh would modulate the innate immune burst at the earliest the early stages of respiratory viral infection. However, we found no evidence increased cholinergic activity during the first week of influenza infection. Although other studies have demonstrated ChAT expression in natural killer cells and myeloid populations(44), we could not unequivocally identify ChAT expression in NK cells, myeloid cells, or alveolar macrophages at any state of infection. Instead we determined that the peak concentration of airway ACh mirrored the kinetics of the airway CD4 T cell population, between 8-10 days after infection. These findings suggested a role in regulating the cellular immune response and/or the transition from active immunity to tissue repair rather than modulating the innate immune burst. Further supporting a role in recovery and repair, cholinergic lymphocyte numbers did not increase during the innate response, but they were retained in the airway and lungs throughout the late stages of infection and long after clinical recovery. Data shown here indicates that cholinergic CD4 T cells are present for at least two months after recovery, well past the time point associated with establishment of the CD4 T cell resident memory population(35, 45). The long-term retention of cholinergic CD4 T cells in the circulation-sequestered niche of the lung, along with their surface phenotype of CD44^hi^CD62U^lo^, lead to speculation that they may also play an important role in the memory T cell response. Since expression of the ChAT-eGFP gene is transient, the ChAT mice used herein cannot be used to determine the full percentage of cells with a cholinergic history. The use of lineage tracer animals will be necessary to fully address these questions.

Despite the clear influx of cholinergic lymphocytes and increased airway choline concentration, we were unable to detect changes in the airway ACh concentration. This is consistent with other studies in the lung(42). The in vivo half-life of ACh is extremely short due to the efficiency of the specific enzymes acetylcholinesterase (AChE) and butyrylcholinesterase (BuChE)(46). To overcome this rapid degradation, cholinergic signaling between cells must take place over a relatively short distance. We found evidence of direct physical contact between Iba1^+^ macrophages and cholinergic lymphocytes in the influenza-infected lung. This contact illustrates a biologic method to ensure proper targeting of the secreted ACh while overcoming the rapid ACh hydrolysis in the pulmonary environment. This direct physical interaction also offers a biological explanation for the presence of cholinergic lymphocytes in a tissue that already possesses two endogenous sources of ACh. The cholinergic vagal nerve can be detected in bronchus-associated lymphoid tissue but it does not enervate the airway. Bronchial epithelium, which lines the airway, is also cholinergic. Our results showed evidence of direct physical interaction between the cholinergic lymphocytes and activated macrophages in all regions of the lungs. In the BALT and peri-bronchial lung regions, multiple sources of ACh may contribute to cholinergic signaling whereas the alveolar space is relatively inaccessible to ACh secreted by bronchial epithelium or the vagal nerve. In addition, although our studies did not determine the primary inducers of ACh secretion by the cholinergic lymphocytes, previous studies have shown that cholinergic lymphocytes produce ACh in response to specific antigen, TLR agonists, neurotransmitters, and cytokines (11-13, 47). As yet we do not know if local neurotransmitter secretion plays any role in lymphocyte ACh production during influenza. For this reason, we have not used the term cholinergic anti-inflammatory pathway, or CAP, since we do not yet what role the brain-immune circuit plays in this process.

Although flow cytometry indicated that CD4 T cells were the dominant cholinergic population in absolute numbers and per-cell ChAT expression, the fluorescent images do not confirm the identity of the cholinergic cell population(s) seen in contact with pulmonary macrophages. The physical contact between cholinergic lymphocytes and activated macrophages throughout the lungs provide evidence for the specialized role of cholinergic lymphocytes as mediators of localized cholinergic signaling and offers a mechanism to avoid deleterious consequences of increased ACh concentration.

The importance of ACh in recovery was indicated by the increased inflammation and aberrant tissue repair noted when Ach synthesis was inhibited. These observations support a necessary, protective role for ACh mediating anti-inflammatory and tissue repair oriented mechanisms in the late stages of infection. ACh as a critical mediator of pulmonary repair following viral illness is consistent with other reports in the literature. ACh increases proliferation of bronchial epithelium, whereas decreased ACh slows proliferation (48). Animals with an endogenous defect in ACh generation exhibit abnormal lung remodeling even in the absence of overt injury (49). Our findings are also supported by studies using different models of acute lung injury (i.e., acid, LPS, *E. coli*), where injury is increased when cholinergic signaling is inhibited and decreased when it is augmented(15, 42, 50).

In addition to inducing epithelial proliferation, ACh mediates changes in macrophage gene expression. The anti-inflammatory activity of ACh binding to the□α7nAchR on macrophages, resulting in diminished NF-κB nuclear translocation and decreased inflammatory cytokine production is well described(39, 51) and has been documented in the lungs(22, 42, 50). Recently, loss of α7nAChR signal transduction was shown to decrease expression of the canonical M2 marker Arginase-1(52). In addition, use of an α7nAchR agonist decreased the LPS-induced inflammatory response and reversed the inflammatory profile, particularly regarding M1 and M2 polarization, while also improving lung function and remodeling in a model of acute lung injury(15). Based on these findings and those in the current study, we hypothesize that ACh produced by cholinergic lymphocytes acts on □□□□□□□□□□macrophages to decrease pro-inflammatory cytokine release and initiate tissue repair during the recovery phase of respiratory viral infection.

In these studies, we used a novel analysis tool to quantify Iba1 expression, relating to pulmonary inflammation. An automated MATLAB algorithm was implemented to accurately isolate the red stain in the Iba1 fluorescent images. The algorithm automatically reads images from a folder sequentially, segments the red stains and computes the amount of red stain present in each of the images. To perform the image segmentation to isolate the red stain, Otsu’s method of thresholding was implemented in order to remove background noise with minimal human bias. The algorithm was automated for computational efficiency and avoids the laborious process of manual segmentation and analysis. Using the algorithm, we investigated Iba1 expression during pulmonary inflammation in different regions of the lungs. The automated algorithm proved to be a valuable tool to quantitate Iba1 expression of pulmonary inflammation, allowing for a quick and accurate analysis of each image with minimal of human bias. The results indicated that Iba1 is involved in the pathogenesis of respiratory viral infection. To the best of our knowledge, this is the first time that Iba1 has been linked to pathology during viral infection. The increased Iba1 expression displayed by infected animals treated with HC3 is also consistent with decreased ACh-induced inflammatory regulation. The histopathological observations in these animals indicate a higher degree of ongoing inflammation when ACh synthesis is pharmacologically disrupted. As an activation marker, Iba1 can be detected on alveolar macrophages as well as myeloid-derived pulmonary macrophages(53). More cells expressed Iba1 in the infected, HC3 treated animals, resulting in larger areas of immunofluorescence. These animals also displayed higher mean staining intensity, indicating that overall Iba1 expression was increased following disruption of ACh synthesis during infection. Although Iba1 has historically been used as an activated macrophage marker(53), the secreted Iba1 protein also acts as an independent inflammatory stimulus. IBA/AIF-1 induces IL-6, TNFα, and CXCL1 (KC) production by pulmonary macrophages and fibroblasts(8, 36). The increased neutrophils in the HC3-treated animals would be consistent with increased pulmonary CXCL1(54).

In the context of our study, we hypothesize that elevated expression of Iba1 following ACh disruption is both indicative of and enhances the ongoing inflammation observed in the lungs. Although the overall role of Iba1 in influenza pathogenesis remains to be elucidated, the overall increased inflammation and delayed recovery in ACh-depleted animals shown here is reminiscent of the response seen in aged animals to respiratory infection(55). Aging is well established as the primary risk factor in murine influenza, human influenza, and currently in the COVID-19 pandemic. Aging also impacts multiple aspects of cholinergic systems beyond expression and cellular distribution of the α7nAchR changes with age, activity, and continued exposure to ACh (reviewed in (56, 57). Aging alters the T lymphocyte response to cholinergic stimulation(58), as well as immune system ACh generation(59, 60). If altered cholinergic capacity plays a role in the diminished immune response or delayed recovery to respiratory infection shown by the elderly, then improving cholinergic function could result in enhanced immune function during aging. Since the elderly are at the greatest risk of death not just from influenza infection but also from the ongoing COVID-19 pandemic, these are questions of the utmost importance. One key feature of age-related immunodeficiency is in the increased basal inflammatory status known as inflammaging(61, 62). Supporting the concept of improving immunity in through cholinergic manipulation, it was first shown over 15 years ago that treatment with the AChE antagonist donepezil decreased inflammatory cytokine message in circulating blood leukocytes of Alzheimer’s Disease patients(63, 64). This study was only recently followed up with the demonstration that inflammatory cytokines TNF, IFN□, IL1b, and IL6 were significantly decreased following six months of donepezil therapy(65). Furthermore, donepezil treatment is associated with decreased overall mortality, including pneumonia-associated mortality (66, 67). As yet, these data are not available for COVID-19 patients. Given the critical unmet need of the elderly for better therapeutic options, extended studies into improved immune function through cholinergic manipulations are of the utmost importance from both a scientific and a world health perspective.

## Summation

ACh has shown to be of extreme importance in recovery from influenza A viral infection. While further studies are needed to elucidate the full mechanistic understanding, ACh has shown to be a critical factor in post-influenza infection recovery. Cholinergic lymphocytes appear in the lungs and airways during the recovery phase of influenza, and are found in direct physical contact with activated macrophages. Artificially decreasing the airway ACh concentration results in extended morbidity, disordered tissue repair, and increased pulmonary inflammation. Together, these results indicate a previously unsuspected role for ACh in mediating pulmonary inflammation and efficient tissue repair during recovery from respiratory viral infection.

## Author Contributions

Conceived and designed the experiments: AH MT DC RF JP.

Performed the experiments: AH CA EH MT DC RF JP.

Analyzed the data: AH AS CA EH MT DC KO RF UG JP

Contributed reagents/materials/analysis tools: AH AS CA MT DC KO RF UG JP.

Wrote the paper: AH AS DC UG JP.

## Competing Interests

The authors declare that no competing interests exist.

